# Deep learning assessment of cultural ecosystem services from social media images

**DOI:** 10.1101/2021.06.23.449176

**Authors:** Ana Sofia Cardoso, Francesco Renna, Ricardo Moreno-Llorca, Domingo Alcaraz-Segura, Siham Tabik, Richard J. Ladle, Ana Sofia Vaz

## Abstract

Crowdsourced social media data has become popular in the assessment of cultural ecosystem services (CES). Advances in deep learning show great potential for the timely assessment of CES at large scales. Here, we describe a procedure for automating the assessment of image elements pertaining to CES from social media. We focus on a binary (natural, human) and a multiclass (posing, species, nature, landscape, human activities, human structures) classification of those elements using two Convolutional Neural Networks (CNNs; VGG16 and ResNet152) with the weights from two large datasets - Places365 and ImageNet -, and our own dataset. We train those CNNs over Flickr and Wikiloc images from the Peneda-Gerês region (Portugal) and evaluate their transferability to wider areas, using Sierra Nevada (Spain) as test. CNNs trained for Peneda-Gerês performed well, with results for the binary classification (F1-score > 80%) exceeding those for the multiclass classification (> 60%). CNNs pre-trained with Places365 and ImageNet data performed significantly better than with our data. Model performance decreased when transferred to Sierra Nevada, but their performances were satisfactory (> 60%). The combination of manual annotations, freely available CNNs and pre-trained local datasets thereby show great relevance to support automated CES assessments from social media.

## 1. INTRODUCTION

Ecosystem services (ES) are defined as the benefits that nature provides to people (MA, 2005). ES include material outputs from ecosystems (e.g., food and raw materials), called provisioning services, as well as the human benefits that derive from ecosystem maintenance and functioning (e.g., water quality and climate regulation), referred as regulating services. ES also include the non-material benefits that humans accrue from nature, known as cultural ecosystem services (CES). These include the benefits that stem from nature-based experiences (e.g., recreation and ecotourism) and intellectual interactions (e.g., aesthetic and heritage values; MA, 2005; Fish et al., 2016a). While initially less studied than other categories of ES (Chan, 2012), CES are seen as increasingly relevant to conservation and sustainable development agendas, as they promote recreation revenues, shape human identity and traditions, and motivate conservation actions (Di Minin et al., 2015). Nevertheless, integrating CES into real-world decision-making is challenging due to the difficulties of operationally defining, quantifying and mapping such intangible and often subjective phenomena (Cheng et al., 2019).

CES are relational (Onofri & Boatto, 2020) and are most frequently evaluated through social surveys. These can produce high quality, contextualized data but necessarily have limited temporal and spatial coverage (Yoshimura & Hiura, 2017) and often suffer from a lack of replication (Bragagnolo et al. 2016). More recently, the emergence of Digital Conservation (van der Wal & Arts, 2015), iEcology (Jarić et al., 2020) and Conservation Culturomics (Ladle et al., 2016) has brought new opportunities to address CES at scale. These disciplinary fields are united by their use of digital (big) data to understand human-nature interactions, to support nature conservation, and/or to promote ecosystem sustainability (van der Wal & Arts, 2015; Toivonen et al., 2019). In contrast to dedicated social surveys, the digital information and data used by these disciplines is frequently produced as a by-product of people’s interactions with the natural environment. For example, through images taken on mobile devices (e.g., smartphones or digital cameras) that are subsequently shared on social media platforms (e.g., through geo-tagged images and messages).

Social media platforms, such as Flickr, contain large amounts of digital data broadly representing people’s interactions with their environment (Di Minin et al., 2015) and, if carefully analysed, can support CES assessments (Toivonen et al. 2019). For instance, content analysis of crowdsourced social media images has allowed researchers to identify visitation patterns, landscape values and human activities in nature (Fu & Rui, 2017; Richards & Tunçer, 2018; Tenkanen et al., 2017). These interactions are frequently amenable to high resolution analysis over large spatial and temporal scales, since the digital content shared on social media platforms often includes georeferenced meta-data and specific time stamps (Gliozzo et al., 2016). Nevertheless, most social media content analyses in the context of CES are based on the manual classification of crowdsourced images or texts shared by social media users (Cheng et al., 2019; Retka et al., 2019). Such manual classification of high volumes of photographic data is time consuming and costly, particularly when it comes to study large geographic areas, time periods and audiences.

The state-of-the art models in the task of image classification, namely deep Convolutional Neural Networks (CNN), have recently been suggested as promising opportunities to assess CES (and other ES; Willcock et al., 2018; Richards & Tunçer, 2018). Deep learning models constitute a class of machine learning algorithms frequently used for automating classification tasks on digital images or videos (e.g., computer vision) and text (e.g., natural language processing; Najafabadi et al., 2015). An example of deep learning models drawing increasing attention from ecology are Convolutional Neural Networks (CNNs; Lusch et al., 2018). CNNs are capable of learning to identify similarities between patterns of information, in a manner that closely resembles a biological brain. Examples of studies that use CNNs include bird individual recognition (Ferreira et al., 2020), plant species identification (James & Bradshaw, 2020) and biodiversity detection (Weinstein, 2018). However, to date the CNN tools that have been used to analyse CES using crowdsourced social media data are not freely available in their full version (Richards & Tunçer, 2018; Gosal et al., 2019), limiting their wider use by researchers, managers, and decision-makers.

This study describes an approach using freely available tools for the automated classification of CES from crowdsourced social media images. Specifically, our study addresses the following questions: (1) Can deep learning CNNs be developed to support an automated classification of crowdsourced social media images to assess CES? (2) Can pre-existent CNN architectures be improved to produce robust and reliable image classifications? and (3) To what extent can pre-trained CNNs be used to classify CES using images from wider geographical contexts? To answer these questions, we first trained different CNN architectures (referred as simply CNNs, hereafter) on social media images (Flickr and Wikiloc) taken for the Peneda-Gerês National Park (Northern Portugal). Then, we optimized those CNNs based on transfer learning (i.e., by pre-training the networks over large image datasets and then re-training them with the Peneda-Gerês data) and data augmentation techniques and compared the robustness of their classification results. Finally, we tested the transferability of those CNNs by applying them to a different image dataset from the Sierra Nevada Biosphere Reserve (Southern Spain).

## 2. METHODS

### 2.1. Methodological framework

Our methodological framework can be divided into three main steps (Fig. 1). First, we manually annotated photos from two image datasets from distinct locations (Peneda-Gerês and Sierra Nevada), previously extracted from the Flickr and Wikiloc platforms. Both datasets were pre-processed to train and validate different automated classification models, based on six CNNs. Second, trained and re-trained six different CNNs using a transfer learning approach and image data from Peneda-Gerês. The classification models were subsequently optimized through data augmentation and changes in model parameters. Finally, we evaluated the performance of the different models by computing a series of classification metrics. The transferability of the best classification models was then tested on the image dataset from Sierra Nevada. All procedures were implemented in Google Colab (https://colab.research.google.com/), using the Keras platform (https://keras.io/) with TensorFlow backend (https://www.tensorflow.org/). These are two of the most used libraries for building and training deep learning models, especially CNNs.

**Fig. 1.**
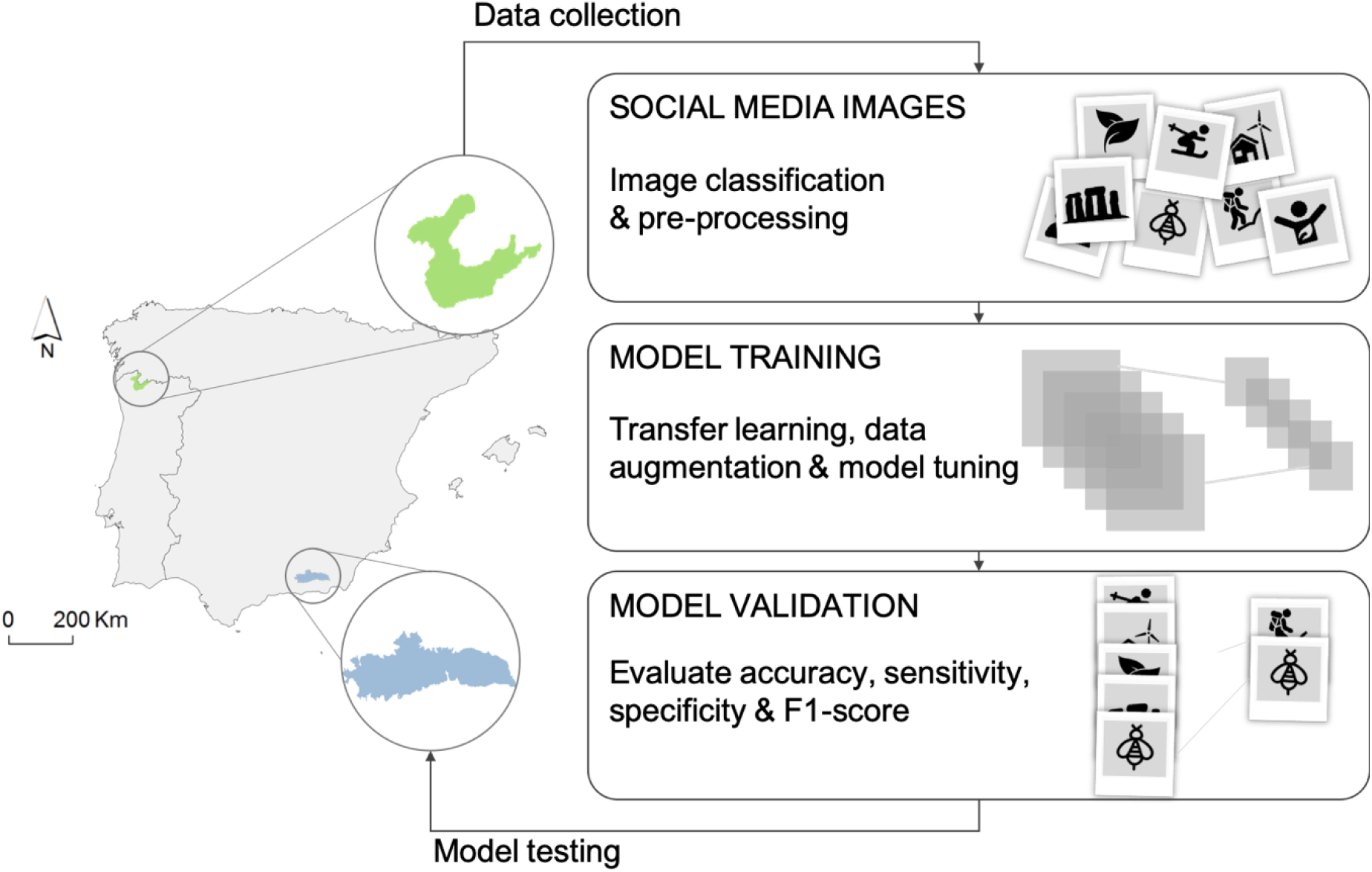
Our methodological framework for developing CNN-based automated classification of CES represented in crowdsourced social media images (see main text for details).

### 2.2. Targeted areas

To provide an automated CES classification from crowdsourced social media images we developed and optimized different CNNs (objectives 1 and 2) using social media images from Peneda-Gerês (our training area). The evaluation of CNN transferability (objective 3) was achieved by using the pre-trained models to assess social media images from Sierra Nevada (our test area; Fig. 1).

Our training area, Peneda-Gerês, is in Northern Portugal and includes the only National Park in the country, and other areas protected under the Natura 2000 network. The climate is Atlantic to sub-Mediterranean and the area holds a rich biodiversity with mountain landscapes characterized by native scrublands, grasslands and *Quercus* woodlands. Peneda-Gerês also contains diverse archaeological and historical sites, including megalithic monuments and signs of Roman occupation. These diverse biophysical and cultural characteristics makes it a very popular area for recreation and other socio-cultural activities (Santarem et al., 2015).

Our test area, Sierra Nevada, is in Southern Spain, and is classified as a UNESCO Biosphere Reserve. It is included in the European Long-Term Socio-Ecological Research Infrastructure and is considered one of the most important biodiversity hotspots in the Mediterranean region, being listed under multiple conservation designations, e.g., Natural and National Parks, Natura 2000 Special Protection Area and Special Area of Conservation. Sierra Nevada supports numerous social-cultural activities with a focus on rural tourism and sport activities (Ros-Candeira et al., 2020).

### 2.3. Crowdsourced social media data

#### 2.3.1. Data mining

Two image datasets were used in this study: for Peneda-Gerês (training area) and Sierra Nevada (test area). Data was collected from two social media platforms: Flickr (https://www.flickr.com/) and Wikiloc (https://www.wikiloc.com/) platforms. For each study area, we collected georeferenced images up to 2018 using the Application Programming Interface (API) of the platforms with Python collection tools. Data protected by social media users’ rights was not considered. Public data containing personal information (including images with recognizable faces) were treated as anonymous. We considered only images taken outdoors; images inside people’s homes were not considered. Likewise, images with irrelevant subjects in the context of CES (e.g., advertisements, pamphlets and drawings) were also excluded. A total of 1778 was considered for Peneda-Gerês and 745 images for Sierra Nevada.

#### 2.3.2. Data labelling

We manually classified each image using a two-step classification process (following Hausmann et al., 2018; Vaz et al., 2019; Moreno-Llorca et al., 2020). In a first level (L1), we used a binary classification, in which each image was labelled as “Natural” or “Human”, depending on whether the image was dominated by natural (Fig. 2a) or human-made (Fig. 2b) elements.

**Fig. 2.**
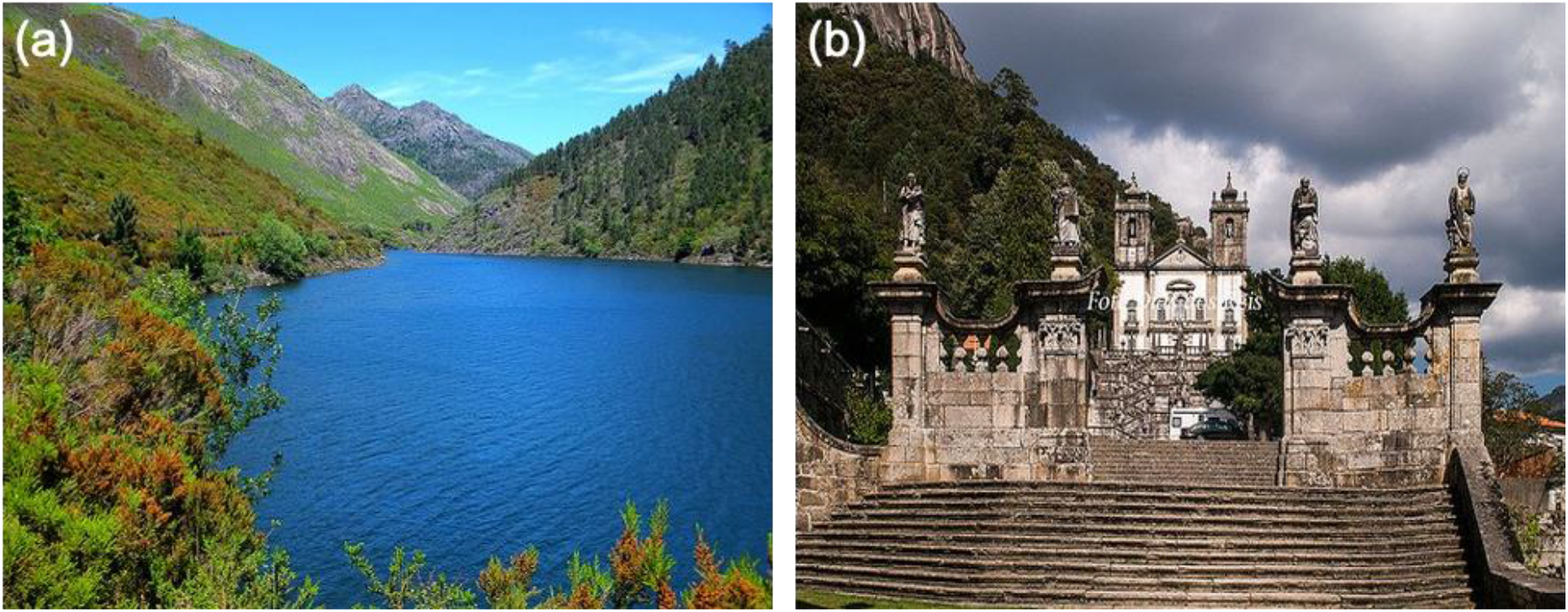
Examples of social media images dominated by natural or human-made elements, being labelled in the first classification level (C1) as: “Natural” (a) or “Human” (b).

In a second level (L2), we used a multiclass classification: “Species”, “Landscape”, “Nature”, “Human activities”, “Human structures” or “Posing” (Fig. 3). “Species” images pertained to close-up shots of animals or plants (Fig. 3a), broadly aligning with the CES of species appreciation (Goodness et al., 2016). “Landscape” images were classified as those depicting ‘wide-open’ shots of nature, often with a visible horizon (Fig. 3b). These were interpreted as representing people’s enjoyment of landscape aesthetics (following Richards & Friess, 2015). “Human activities” include photos of people engaging in recreational activities (Richards & Friess, 2015). For instance, photos of sporting activities such as skiing or cycling (Fig. 3c). The “Human structures” category included photos where human-made structures dominated the photo, e.g. historical monuments and churches (Fig. 3d). These were interpreted as representing CES relating to cultural heritage and spiritual enrichment (Blicharska et al., 2017). “Posing” category refers to photos of people looking at the camera, with recognizable faces (Fig. 3e). Such an interaction with the photographer is representative of CES relating to social enjoyment and sense of identity (Riechers et al., 2016). Finally, the “Nature” category relates to images that include natural elements but with no particular feature (such as species) and taken at intermediate ranges (differing from the more distant perspectives used in landscape photos, Fig. 3f). This category was interpreted as representing a general appreciation of nature by people (following Richards & Friess, 2015). Details on the data collection and manual classification processes can be found in the previous studies from Vaz et al. (2019), Ros-Candeira et al. (2020) and Moreno-Llorca et al. (2020).

**Fig. 3.**
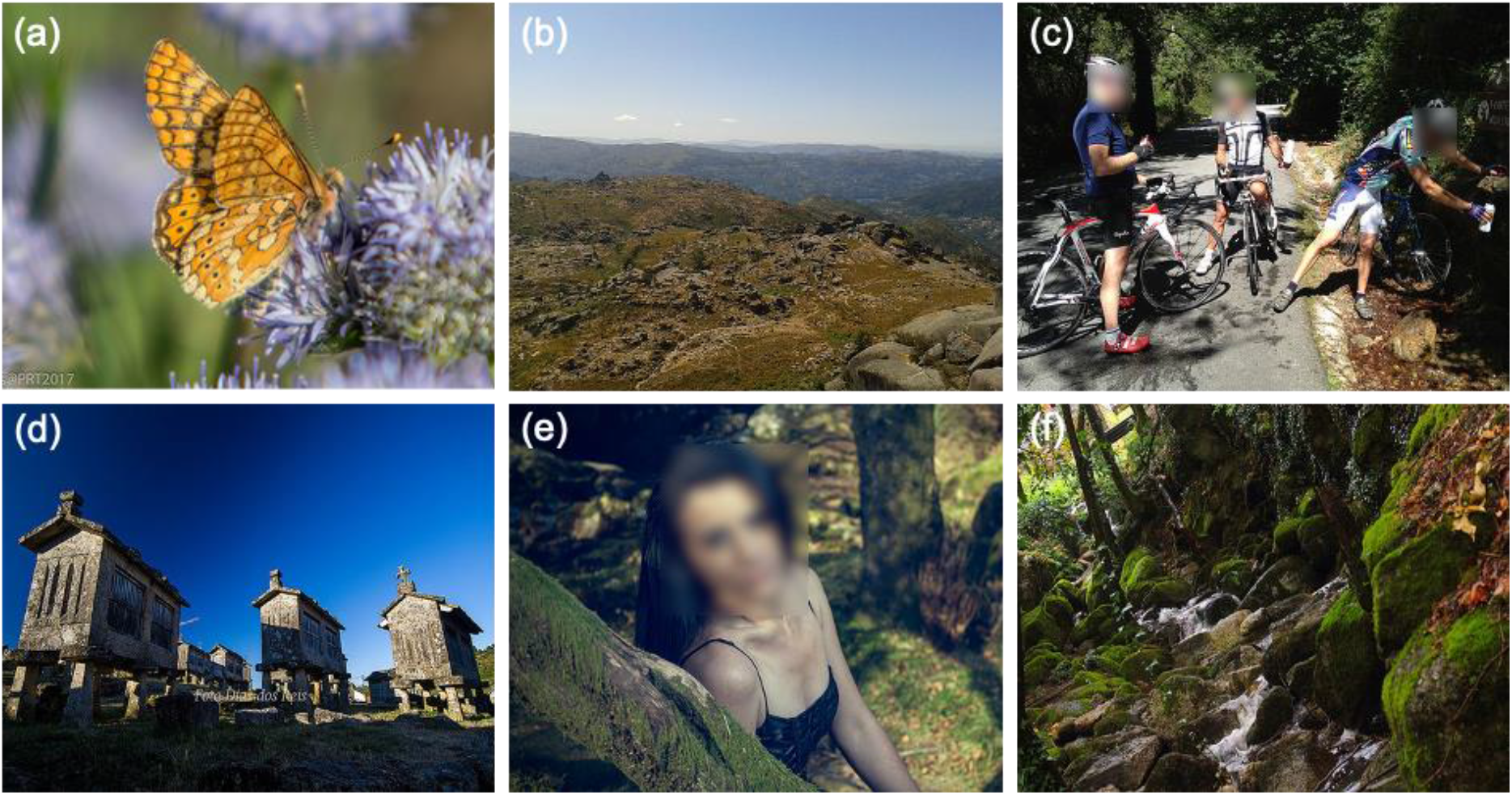
Examples of social media images being labelled in the second classification level (C2), as “Species” (a), “Landscape” (b), “Human activities” (c), “Human structures” (d), “Posing” (e) and “Nature” (f).

From Peneda-Gerês (our training area), the proportion of images in the classes “Natural” or “Human” was 48% and 52%, respectively. From Sierra Nevada (our test area), 62% of all images belonged to the “Natural” class and 34% belonged to the “Human” class. For the L2 classification, photos from Peneda-Gerês were classified as 35% “Landscape”, 28% “Human structures”, 13% “Nature”, 12% “Species”, 7% “Posing” (7%), and 5% “Human activities”. In Sierra Nevada, most (38%) images belonged to the “Human structures” class, 19% to “Landscape”, 11% to “Human activities” and “Posing”, and 10% to “Nature” and “Species” (Fig. 4).

**Fig. 4.**
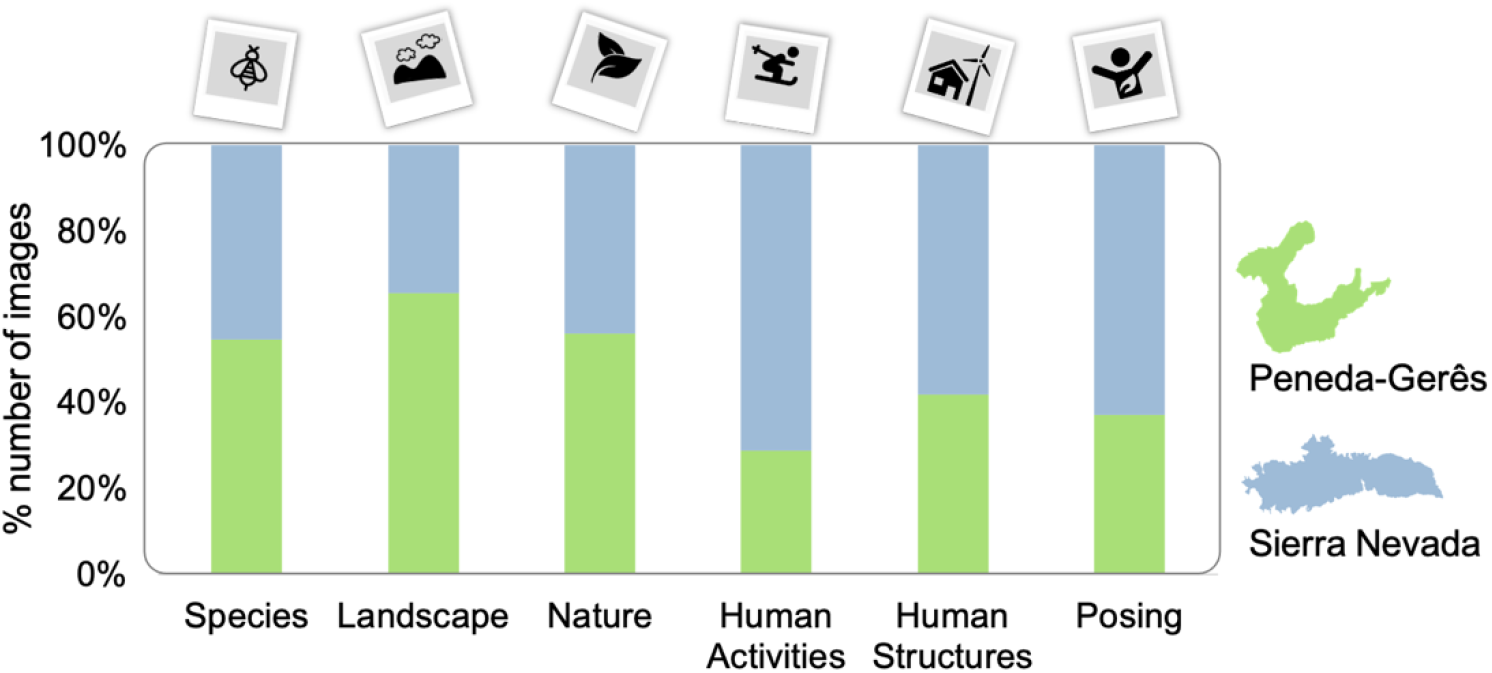
Proportion of social media images assigned manually to each class of cultural ecosystem services in Peneda-Gerês and Sierra Nevada.

### 2.4. Model training

To efficiently train and test CNNs, a series of pre-processing procedures normally need to be applied (Na & Fox, 2020). Specifically, before model training all images were resized to the same resolution (473 × 372 pixels) by taking into account the mean resolution of the set, and then normalized to the [0,1] range.

#### 2.4.1. Convolutional Neural Networks (CNN)

To automate classification of the image dataset, two of the most influential CNNs were considered: VGG16 (Very Deep Convolutional Network) and ResNet152 (Residual Neural Network). These CNNs have high performance and benefit from a vast amount of pre-trained weights freely available online (Dramsch & Lüthje, 2018).

VGG16 is considered very effective for large-scale image recognition and classification tasks (Simonyan & Zisserman 2014). During training, each image passes through a stack of convolutional layers, where two different types of filters are applied: i) a 3×3 filter, which captures the notion of left/right, up/down and centre, and; ii) a 1×1 filter, which provides a linear transformation of the input channels. Following filtering, max pooling is applied to the images using 2×2 kernels, followed by three fully connected layers and a SoftMax layer, that provides a probability distribution for each class.

ResNet152 makes use of residual learning units. These units avoid the problem of vanishing gradients (He et al., 2016), because activations from a previous layer are reused until the next layer learns its weights. ResNet architectures are often implemented using double or triple layer nonlinear jumps with batch normalization, meaning that each layer is connected to the next one and to the layers 2-3 jumps away (He et al., 2016). During training, the model keeps layers that are contributing to improvements in the classification performance and discards layers compromising effectiveness.

In both CNNs, we used the ADAM algorithm as optimizer and a batch size of 10 (Kingma & Ba, 2014). An early stop approach was implemented, with a patience value of 16, meaning that if there was no decrease in the model loss (binary cross entropy) for more than 16 consecutive epochs we would stop training. For both L1 and L2 classifications, the best learning rates for VGG16 and Resnet152 were 10^−6^ and 10^−4^, respectively, which were chosen from empirical trials over 100 epochs. Both VGG16 and ResNet152 models showed no improvements in model accuracy and loss after 50 epochs. Details on CNN parameterization and implementation can be found in the Supporting Information.

#### 2.4.2. Training and validation datasets

Training an image classification model (in our case, a CNN) requires a training and a validation dataset. The training dataset is the subset of images that the neural network will use to learn how to classify the input images into different classes (in our case, representations of different CES), by adapting the weights in its layers. The validation dataset is an independent subset of images used to estimate the error of the model during training (accuracy and loss) and to assess the learning progress of the neural network. The validation dataset indicates if the model is overfitting the training data and not learning features that are essential for the recognition of different classes.

We used the k-fold cross-validation by partitioning the Peneda-Gerês dataset into *k* equal sized random subsets. One of these subsets was retained to evaluate model performance, while the remaining *k*-1 subsets are used for training. This procedure is repeated so that each subset is used only once for validating the model. In this study, a 5-fold-cross validation method was adopted, meaning that the original dataset was randomly partitioned into 5 equal sized subsets. During the training of the model, in each of the 5 folds, 90% of the data of the 4 subsets was used for training the model. The remaining 10% was used for model validation, allowing us to determine the best training parameters and to evaluate the performance of the model.

To further increase the utility of our training dataset, we used a data augmentation procedure to artificially increase the dataset size by applying random transformations to the existing set of images. In our training dataset, five random transformations per image were considered using the data generator available in Keras (Chollet, 2015). Specifically, each image was randomly rotated (from 0 to 40°) and zoomed (zoom range of 0.2). We further applied horizontal flip as well as width (width shift range of 0.2) and height shift (height shift range of 0.2) to the training images. Both original (before data augmentation) and modified (after data augmentation) images were used, resulting in a final dataset of 8532 or 8538 images, depending on the fold.

#### 2.4.3. Transfer learning

To improve the generalization of the CNNs and to avoid possible overfitting due to the small size of our training data, we initialized both VGG16 and ResNet152 models using the weights of pre-trained networks (transfer learning). This has the added advantage of using network architectures which have already learned features (such as colour or texture) that are relevant to distinguish multiple objects and that can be useful to distinguish our CES classes.

We tested weights of freely available networks pre-trained on the “Places365” (https://github.com/CSAILVision/places365) and “ImageNet” (http://www.image-net.org/) dataset. The “Places365” dataset is part of the larger “Places” dataset, containing around 1.8 million scene images, labelled with 365 scene semantic categories with similar elements to those tackled by this study (Zhou et al., 2017). The ImageNet database is a large-scale hierarchical image database with multiple applications, comprising more than 14 million cleanly annotated images spread over 21 000 categories and providing representative and diverse coverage of the image world (Mettes et al., 2016). To evaluate the added value of the transfer learning procedure, we considered both VGG16 and ResNet152 with weights obtained from scratch (referred to as “OwnWeights”, hereafter). In total, we considered six CNN models from the combination of the two architectures (VGG16 and ResNet152) with three weight sets (Places365, ImageNet, OwnWeights).

Our aim was to achieve a balance between CNNs capacity to generalize and their adaptation (specificity) to the task of classifying CES in images from our training site. To achieve this balance the first layers of the networks were not updated during training when using transfer learning. We did this because these layers usually extract simple geometrical structures from the images which are applicable to different classification tasks. Thus, the weights of the first layers were those obtained when pre-training with Places365 or ImageNet datasets. To ensure specificity, three fully connected layers in VGG16 and one fully connected layer on ResNet152 were re-trained (fine-tuned) using our training dataset. Moreover, an additional dense layer with 128 units and a rectifier linear unit activation function was included before the output layer, to allow better adaptation of the pre-trained architectures to the classification task. Finally, the output layer was modified to fit a binary classification (with 2 units) that suited our L1 classification (“Natural” and “Human”), as well as our six classes, with 6 units that match the L2 classification (“Species”, “Landscape”, “Nature”, “Human activities”, “Human structures” and “Posing”).

### 2.5. Performance metrics

The performance of our models was evaluated on overall accuracy (ACC), sensitivity (recall or True Negative Rate: TNR), specificity (precision or True Negative Rate: TNR) and F1-score (F1) (see Table 1a for details). In the L1 classification, “Human” images were assumed as positives, and “Natural” images as negatives (Table 1b).

**Table 1a.** Evaluation metrics considered in the model evaluation, with respective calculation, description and example from L1 classification.

| Metric | Calculation | Description | Example |
| --- | --- | --- | --- |
| Accuracy | $ACC = \frac{TP+TN}{TP+TN+FP+FN}$ | Ratio of correctly predicted labels by the model to the total images. | Shows the level of correctly classified images as “Human” and as “Natural” by the model. |
| Sensitivity<br>(or recall) | $TPR = \frac{TP}{TP+FN}$ | Ratio of correctly predicted labels by the model to all images in actual label. | Indicates the proportion of correctly classified images as “Human” by the model, in relation to all actual “Human” images. I.e., *Of all images that were actually “Human”, how many did the model predict as “Human”?* |
| Specificity<br>(or precision) | $TNR = \frac{TP}{TP+FP}$ | Ratio of correctly predicted labels by the model to the total of predicted positive labels. | Indicates the proportion of all classified images as “Human” by the model, in relation to all predicted “Human” images. I.e., *Of all images labelled as “Human” by the model, how many were actually “Human”?* |
| F1-score | $F_1 = \frac{2TP}{2TP+FP+FN}$ | Computes the harmonic mean of the specificity and sensitivity. | Shows the level of correctly and incorrectly classified images as “Human” and as “Natural” by the model. |

**Table 1b.** Example of a confusion matrix obtained based on the L1 classes “Human” and “Natural”.

|  |  | Predicted label as classified by the model |  |
| --- | --- | --- | --- |
|  |  | “Human” | “Natural” (not “Human”) |
| Actual label as<br>manually classified | “Human” | True Positive (TP) | False Negative (FN) |
|  | “Natural” (not<br>“Human”) | False Positive (FP) | True Negative (TN) |

For the L1 classification, the previous classification metrics were computed as the mean of the performance metrics obtained over the 5 different folds. For the L2 classification, for each fold, the performance metrics were calculated following a “one-versus-all” definition, i.e., by considering as positives the images of each label/class, while considering as negatives all the images from the remaining labels/classes. Metrics were computed via the macro-average; the unweighted mean of the metrics obtained for each class (Thorat et al., 2020). We used a paired samples t-test, with a confidence interval of 0.05 (Hsu & Lachenbruch, 2005) to test for significant differences in classification metrics between each pair of the six CNN models.

### 2.6. Model testing

The best performing models for classifying the Peneda-Gerês dataset were used to classify the image dataset from Sierra Nevada. Model performance on the new dataset was evaluated using the metrics previously described, with the aim of assessing whether CNN models trained and re-trained in Peneda-Gerês could effectively classify CES in images from Sierra Nevada (objective 3).

## 3. RESULTS

### 3.1. CNN performance

The six CNN models performed differently for each image classification level (L1 and L2; Table 2). Overall, the ResNet152-ImageNet performed best for both classifications (followed by VGG16-Places365) in terms of mean accuracy, sensitivity, and F1-score (see Supplementary material for details on confusion matrices). The models weighted from our own dataset always generated the least satisfactory results.

**Table 2.** Classification metrics obtained for the six models: both VGG16 and ResNet152 CNNs were re-trained after their weights were initialized from pre-training over the Places365 and ImageNet, or they were trained from scratch with our data (mean ± standard deviation of the five folds). Results are shown for both L1 and L2 classifications. Bold values highlight the best performance results for each classification level and metric.

| Models / Weights | L1 Classification |  | L2 Classification |  |
| --- | --- | --- | --- | --- |
|  | VGG16 | ResNet152 | VGG16 | ResNet152 |
| *Accuracy* |  |  |  |  |
| Places365 | **87.01 ± 2.55** | 86.00 ± 2.86 | **74.07 ± 1.23** | 76.88 ± 2.59 |
| ImageNet | 86.11 ± 2.00 | **87.18 ± 1.75** | 73.73 ± 1.57 | **76.89 ± 1.79** |
| OwnWeights | 72.10 ± 3.94 | 76.43 ± 1.02 | 57.03 ± 3.67 | 61.98 ± 2.25 |
| *Sensitivity* |  |  |  |  |
| Places365 | **88.48 ± 2.47** | 83.40 ± 6.33 | **64.35 ± 2.17** | 67.92 ± 2.54 |
| ImageNet | 86.71 ± 2.85 | **86.78 ± 2.97** | 63.85 ± 2.80 | **68.09 ± 2.57** |
| OwnWeights | 71.54 ± 2.06 | 76.55 ± 4.36 | 42.11 ± 2.34 | 49.69 ± 1.44 |
| *Specificity* |  |  |  |  |
| Places365 | **85.54 ± 3.75** | **88.46 ± 3.03** | **94.32 ± 0.31** | **94.90 ± 0.55** |
| ImageNet | 85.46 ± 1.58 | 87.52 ± 1.24 | 94.30 ± 0.37 | 94.88 ± 0.28 |
| OwnWeights | 72.82 ± 9.35 | 76.39 ± 4.05 | 90.59 ± 0.68 | 91.66 ± 0.45 |
| *F1-Score* |  |  |  |  |
| Places365 | **87.53 ± 2.79** | 85.89 ± 3.60 | **64.84 ± 2.00** | 68.86 ± 2.52 |
| ImageNet | 86.53 ± 2.60 | **87.44 ± 2.39** | 63.92 ± 2.80 | **68.92 ± 2.70** |
| OwnWeights | 72.69 ± 3.51 | 77.23 ± 2.34 | 41.30 ± 1.87 | 49.64 ± 1.55 |

For both L1 and L2 classifications, differences among the six CNN models for all performance metrics were generally not significant (p > 0.05) (see Table S2 for full results). For L1 models, significant differences were only found for sensitivity and F1-score, with ResNet152-OwnWeights performing better than VGG16-OwnWeights (p = 0.03 and p = 0.05, respectively). In this classification, model performance was lower on images dominated by water or rocks; these were usually manually labelled by us as “Natural” and classified by the model as “Human” (see Fig. S1).

For the L2 classification, the ResNet152-ImageNet and ResNet152-OwnWeights models were significantly more accurate than VGG16-ImageNet (p = 0.03) and VGG15-OwnWeights (p = 0.05), respectively. The same pattern was repeated for sensitivity and specificity metrics, in which ResNet152-OwnWeights performed better than VGG16-OwnWeights (p = 0.01 and p = 0.03, respectively). The F1-score also revealed significant differences. ResNet152-Places365 and ResNet152-OwnWeights performed better than VGG16-Places365 (p = 0.05) and VGG16-OwnWeights (p = 0.01), respectively. Classification models were less capable of distinguishing “Landscape” images from “Human activities” or “Human structures” and were also poor at distinguishing “Nature” images from “Species” (Fig. S2).

### 3.2. CNN transferability and generalization

The CNN model (ResNet152-ImageNet) pre-trained over the ImageNet dataset and re-trained with the Peneda-Gerês dataset was less effective when applied to the Sierra Nevada images. Nevertheless, for the L1 classification the model performance was still relatively good, with an accuracy of 72.89%, sensitivity of 85.36%, specificity of 65.38% and F1-score of 70.29% (Fig. 5a). Model classification failed most often on images dominated by snow, which were manually labelled as “Natural” but automatically classified as “Human” (see Fig. S3).

**Fig. 5.**
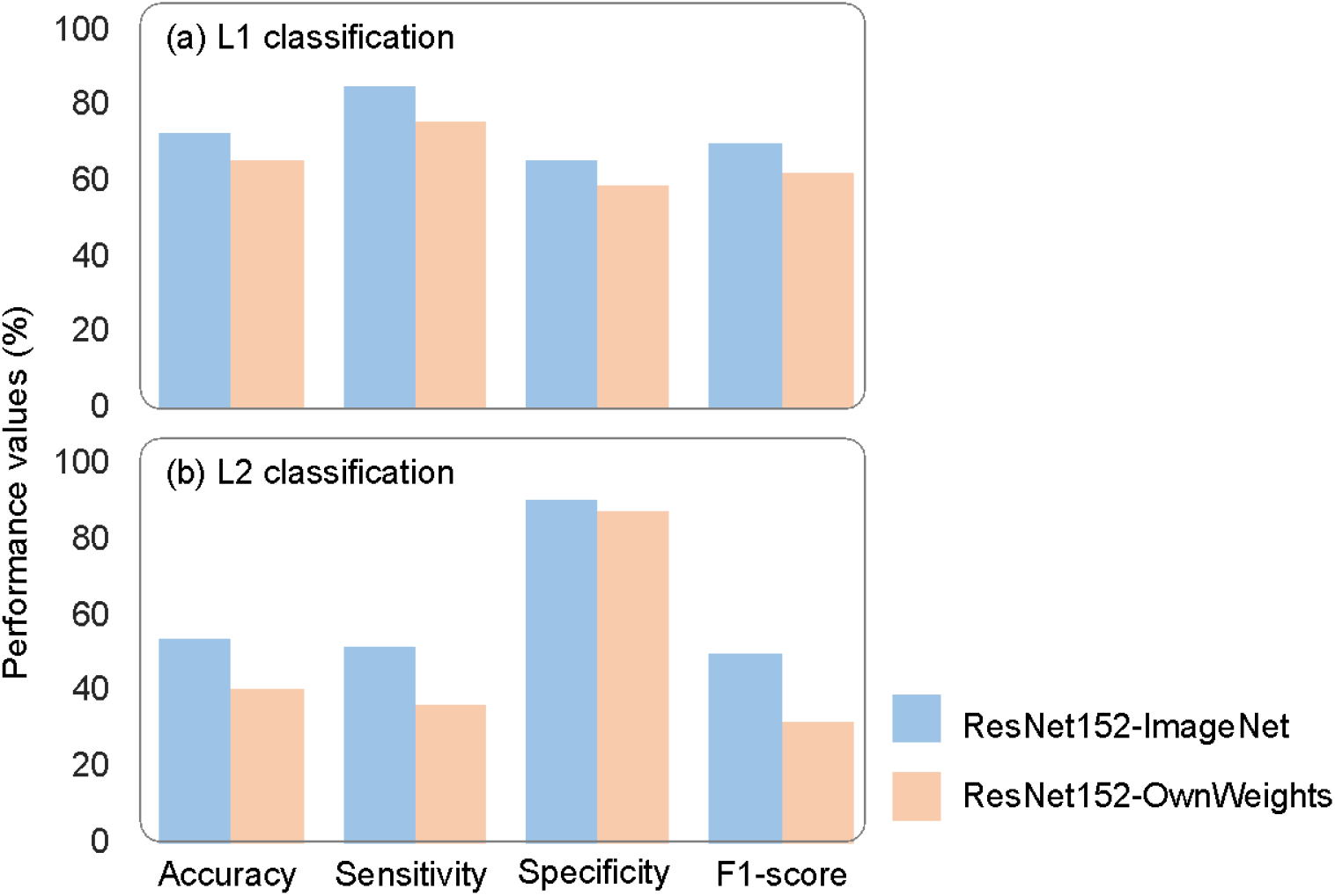
Classification metrics obtained for the ResNet152-ImageNet models, pre-trained over the ImageNet dataset and re-trained with the Peneda-Gerês dataset, applied to the Sierra Nevada dataset. Results from the ResNet152 built with the weights from our own dataset is also exhibited as control.

For the L2 classification, model performance in the Sierra Nevada images was rather poor, with values below 60% for accuracy, sensitivity, and F1-score. However, specificity reached 91% (Fig. 5b). Image classification mostly failed when trying to distinguish “Landscape” from “Nature” images, and “Posing” from “Human activities” images (see Fig. S4).

## 4. DISCUSSION

The use of deep learning tools is becoming increasingly popular in ecology (Wäldchen & Mäder 2018; Christin et al., 2019; Weinstein, 2018). Particularly prominent is the adoption of CNNs to assist on the identification of taxa from images obtained, e.g., from citizen science platforms (e.g., Terry et al., 2019), camera traps (e.g., Sadegh Norouzzadeh et al., 2020), and even feeding structures (e.g., Ferreira et al., 2020). In the research of CES, deep learning is also showing great potential, helping on the identification of human-nature interactions from crowdsourced image datasets shared in online social media (Gosal & Ziv 2020; Jarić et al. 2020). Here, we applied six different CNN models to support the classification of crowdsourced social media images, based on common outdoor features underlying CES assessments. Our approach compared the performance of two state-of-the art CNNs (ResNet152 and VGG16) built with the weights from three image datasets (Places365, ImageNet, and our own training dataset). The models were trained and re-trained on social media images taken in the Peneda-Gerês protected area. The capacity of the models to successfully classify images from other areas (transferability) was tested by applying them to images from the geographically proximate but biophysically and culturally distinct Sierra Nevada Biosphere Reserve.

In general, the six models trained or re-trained for Peneda-Gerês performed well (cf. Table 2). This is in line with similar studies that have achieved satisfactory performances when classifying nature-based images taken by the public, either through freely available platforms, such as TensorFlow (e.g., Terry et al., 2019), or payable ones, such as Google Vision (e.g., Gosal & Ziv, 2020). These results strongly support the contention that deep learning CNNs can successfully and automatically classify crowdsourced social media images to assess CES (Objective 1). Nevertheless, our CNNs were better able to distinguish a simple dichotomy of “Natural” from “Human” images (i.e., our L1 classification) than our more complex classification of CES (i.e., our L2 classification). Nevertheless, the L2 classification showed higher specificity values than the L1 classification, which tend to highlight differences between the true positive (one class) and the true negatives (several remaining classes) (Silva & Villela, 2020). These findings were anticipated, given that the increasing complexity of model tasks and output nodes required to tackle a multiclass classification inevitably affects the reliability of the results (Li et al., 2018). In our dataset, certain CES classes (e.g., “Landscape” and “Nature”; cf. Fig. 4), often contain many common colour features, textures and patterns (e.g., pertaining to sky, sea, and vegetation), constituting a great challenge for the CNN models (Srivastava et al., 2018). Additionally, some images can be placed in more than one CES class (e.g., posing in a landscape setting or in front of an infrastructure), emphasizing the difficulty in capturing relational issues often associated to CES assessment (e.g., Fish et al. 2016). A size-based rule can be used in the future to solve this obstacle whereby it is designated a single CES based on the proportion of the image that different elements cover.

Overall, the CNN models pre-trained with the ImageNet and Places365 and re-trained over the Peneda-Gerês dataset showed significantly better performances compared to models trained directly with our data, regardless of the classification task (binary and multiclass; cf. Table 2). This verifies that transfer learning can significantly improve image classifications to produce more robust and reliable CNN models (our objective 2). ResNet was the most satisfactory architecture compared to VGG16, performing significantly better at the complex tasks associated with multiclass classification (cf. Table 2). As has been previously noted, the ResNet architecture is particularly robust compared to other architectures, such as AlexNet or VGG (He et al., 2016). In our training dataset, the ResNet pre-trained with the ImageNet weights outperformed the other models (cf. Table 2). This was perhaps surprising, considering that images from the Places365 are almost exclusively focused on landscapes and the wild environment (Seresinhe et al., 2017). Nevertheless, ImageNet has a larger and more diverse image dataset (ca. 14 million images), compared to Places365 (ca. 1.8 million images), which can improve model performance.

When we applied the pre-trained ResNet-ImageNet model to a different area (Sierra Nevada), as anticipated we observed a decrease in model performance (cf. Fig. 5). However, the decrease in performance was relatively minor for accuracy and sensitivity (73%-85%), suggesting that a pre-trained model could still be useful to classify CES from different geographical and cultural contexts (our Objective 3). The loss of model efficacy probably reflects its failure to capture distinctive features (including colours, textures, and patterns) of the new biophysical and cultural environment (Ying, 2019). The model performance declined further for the multiclass classification (< 60% accuracy), due to its inability to distinguish “Landscape” versus “Nature” images, as well “Posing” versus “Human activities” images. In the images from Sierra Nevada, cold and neutral colours (combining white, grey, and blue) tended to dominate, while in Peneda-Gerês, warm and cold colours (combining green, blue and brown), are more prominent. For example, snow is more characteristic from Sierra Nevada but is rarely observed in Peneda-Gerês. Finally, it is important to note that the training area and the test area are relatively geographically and culturally proximate and that the well-known biogeographical pattern of distance decay of biotic similarity (Nekola & White 1999) would make it increasingly challenging to apply pre-trained models to more geographically distant areas.

The use of CNN models to assess CES necessarily requires a degree of fine-tuning and optimization. Nevertheless, the impressive results obtained in this study demonstrate that a combination of manual classifications, freely available CNNs and pre-trained local datasets can be extremely effective to evaluate CES over large regions, especially if these regions are biophysically and culturally homogenous. As the field of deep learning progresses, new tools and solutions will be developed that allow CNNs to classify broader arrays of images over greater geographical scales. In this context, the free availability (and continued growth) of large image datasets, such as the ones used here (ImageNet and Places365), is of the utmost importance (cf. Ferreira et al. 2020).

Moving forward, deep learning analysis of CES could benefit from incorporating other forms of data on human-nature interactions in digital space. For example, most users of file sharing platforms, such as Flickr and Instagram, provide contextual information about their images in the form of tags and captions (e.g., “Contemplating at a beautiful landscape”, “What species is this?”). Integrating other data sources besides image content, may therefore provide a possible way forward to improve the generalization of deep learning tools in the scope of CES analysis (Cai & Xia, 2015; Ladle et al. 2017). More generally, deep learning tools may be equally effective (or even more effective) for evaluating other classifications of the benefits that nature provides to people, especially those focused on practices of human engagement with the natural world (cf. Jepson et al. 2017).

We hope our work will stimulate others to share their datasets, for instance through a global Cultural Ecosystem Services Assessment (CESA) database, in order to further expand the use of deep learning for studying human-nature interactions. Having large datasets will: i) allow better optimization of CNNs; ii) strengthen existing network architectures (or allowing new ones to be built), and; iii) provide data to fine-tuning the hyper-parameters settings, improving results. Moreover, sharing codes and having freely available deep learning tools (such as TensorFlow through Google Colab) with user-friendly programming languages (e.g. Python) under minimum yet robust computational resources can aid researchers, managers, and other stakeholders in their real-world decisions about the management of the natural world and the people that benefit from it.

## ACKNOWLEDGEMENTS

This study acknowledges support from Ministerio de Ciencia, Innovación y Universidades (Spain) through the 2018 Juan de la Cierva-Formación program (contract reference FJC2018-038131-I). This paper contributes to the GEO BON working group on Ecosystem Services.

## DATA AVAILABILITY

All code used in model training and test is available at: https://github.com/anasccardoso/Deep-learning-for-cultural-ecosystem-services. The dataset of “Sierra Nevada” is available at https://figshare.com/articles/Dataset_SN_csv/8943509/2. The dataset of “Peneda-Gerês” will be made available in a public repository (https://zenodo.org/) after acceptance.

