## Supplemental Material for "Deep learning assessment of cultural ecosystem services from social media images"

### SUPPLEMENTARY MATERIAL

#### CNN architectures

VGG16 results from the connection of different combinations of convolutional and pooling layers, followed by three fully connected layers also known as dense layers. This architecture has 16 learnable layers, 13 convolutional layers and 3 fully connected layers. Pooling layers, unlike the remaining layers, are not learnable (Khan et al., 2020). They are actually reduction layers used to increase the abstraction level of the features extracted by the convolutional layers. The stack of convolution and pooling layers are used for extracting increasingly higher-level features from the input images. Lower level layers extract lower level features such as edges, blobs and one colour, to complex features such as complex combinations of different edges, colours and textures. The last three fully connected layers work as a classifier, which main function consists in converting the higher-level features into predictions.

ReNest152 is a more modern architecture that was designed by Microsoft Asia. The main problem of the architectures that precede ResNet152, such as VGG16, is the pooling layers, which eliminate important information extracted along the network. The ResNet152 architecture alleviates this limitation using the residual module, which is the building block of this network (He et al., 2016). The residual module combines the intact input information with the residual information.

### CNN parameterization and implementation

Regarding the optimizers and training parameters of the CNN models, Adam was the selected optimizer, an algorithm for first-order gradient-based optimization of stochastic objective functions, since it is one of the most used optimizers in the scope of deep learning and was indeed the most suitable for the classification models under study (Kingma & Ba, 2014). For batch size, an hyperparameter of gradient descent responsible for controlling the number of training samples to be considered before the model's internal parameters are updated (Brownlee, 2016), it was chosen a mini batch size of 10. This value was selected mainly because of the small size of our data and, especially, due to limitations on available memory usage. At an initial stage, for the six models, it was considered the Keras default learning rate (0.001), as well as 100 epochs. After empirical trials, the learning rates that led to a better model performance were $10^{-6}$ for VGG16 and $10^{-4}$ for ResNet152. Both VGG16 and ResNet152 models showed no improvements in model accuracy and loss after 50 epochs.

An early stop approach was also considered, which is a method used to prevent overfitting when training with an interactive algorithm, such as gradient descent, being widely used in the most diverse fields of deep learning due to its simplicity of understanding and implementation (Nielsen et al., 2017). To achieve that, it was generated a validation set, setting aside 10% of the training data, in order to use the validation loss as the stopping criteria. In other words, at the end of each epoch, the validation loss is computed on the validation data, terminating when it stops improving (i.e., when it stops decreasing). A patience value of 16 was also established, adding a delay to the trigger in terms of the number of epochs (16) on which we expect to perceive no improvement.

**Confusion matrices**

This section includes the confusion matrices of the ReNet152 models, that were obtained for both L1 and L2 classifications.

**Table S1a.** Confusion matrix of ResNet152 with Places365 weights in the L1 classification. 0 – Nature, Human – 1.

| | | Predicted Values | |
|---|---|---|---|
| | | 0 | 1 |
| Actual Values | 0 | 152 | 20 |
| | 1 | 30 | 154 |

**Table S1b.** Confusion matrix of ResNet152 with ImageNet weights in the L1 classification. 0 – Nature, Human – 1.

| | | Predicted Values | |
|---|---|---|---|
| | | 0 | 1 |
| Actual Values | 0 | 150 | 21 |
| | 1 | 24 | 160 |

**Table S1c.** Confusion matrix of ResNet152 with OwnWeights in the L1 classification. 0 – Nature, Human – 1.

| | | Predicted Values | |
|---|---|---|---|
| | | 0 | 1 |
| Actual Values | 0 | 129 | 40 |

| | | |
|---|---|---|
| 1 | 44 | 143 |

**Table S1d.** Confusion matrix of ResNet152 with Places365 weights in the L2 classification. 0 – Species, 1 – Landscape, 2 – Nature, 3 – Human activities, 4 – Human structures, 5 – Posing.

| | | Predicted Values | | | | | |
|---|---|---|---|---|---|---|---|
| | | 0 | 1 | 2 | 3 | 4 | 5 |
| Actual Values | 0 | 31 | 3 | 3 | 1 | 4 | 1 |
| | 1 | 3 | 105 | 6 | 1 | 10 | 2 |
| | 2 | 4 | 6 | 34 | 1 | 3 | 0 |
| | 3 | 1 | 6 | 1 | 5 | 2 | 2 |
| | 4 | 1 | 9 | 4 | 1 | 84 | 1 |
| | 5 | 1 | 2 | 0 | 2 | 2 | 16 |

**Table S1e.** Confusion matrix of ResNet152 with ImageNet weights in the L2 classification. 0 – Species, 1 – Landscape, 2 – Nature, 3 – Human activities, 4 – Human structures, 5 – Posing.

| | | Predicted Values | | | | | |
|---|---|---|---|---|---|---|---|
| | | 0 | 1 | 2 | 3 | 4 | 5 |
| Actual Values | 0 | 30 | 3 | 5 | 1 | 2 | 1 |
| | 1 | 2 | 107 | 6 | 1 | 7 | 1 |
| | 2 | 1 | 6 | 38 | 0 | 1 | 0 |
| | 3 | 1 | 5 | 2 | 4 | 1 | 3 |
| | 4 | 1 | 13 | 6 | 1 | 78 | 1 |
| | 5 | 1 | 3 | 1 | 2 | 2 | 16 |

**Table S1f.** Confusion matrix of ResNet152 with OwnWeights in the L2 classification. 0 – Species, 1 – Landscape, 2 – Nature, 3 – Human activities, 4 – Human structures, 5 – Posing.

| | | Predicted Values | | | | | |
|---|---|---|---|---|---|---|---|
| | | 0 | 1 | 2 | 3 | 4 | 5 |
| Actual Values | 0 | 22 | 5 | 5 | 2 | 6 | 3 |
| | 1 | 2 | 99 | 5 | 3 | 13 | 3 |
| | 2 | 8 | 8 | 23 | 1 | 7 | 1 |
| | 3 | 2 | 8 | 2 | 1 | 2 | 1 |
| | 4 | 6 | 16 | 9 | 1 | 64 | 3 |
| | 5 | 2 | 5 | 1 | 1 | 5 | 10 |

**Table S2.** P-values from the paired t-test for the six models: both VGG16 and ResNet152 CNNs combined with three different weights from Places365, ImageNet, and from our own training dataset. Results are shown for both L1 and L2 classifications. Bold values highlight the statistically significant results.

| Models/Weights pairs | p-value | | | |
|---|---|---|---|---|
| *L1 classification* | *Accuracy* | *Sensitivity* | *Specificity* | *F1-score* |
| VGG16 vs ResNet152 with Places365 | 0.185 | 0.088 | 0.241 | 0.084 |
| VGG16 vs ResNet152 with ImageNet | 0.303 | 0.951 | 0.082 | 0.381 |
| VGG16 vs ResNet152 with OwnWeights | 0.083 | **0.031** | 0.303 | **0.049** |
| VGG16 with Places365 vs VGG16 with ImageNet | 0.516 | 0.131 | 0.966 | 0.453 |
| ResNet152 with Places365 vs VGG16 with ImageNet | 0.287 | 0.159 | 0.517 | 0.197 |
| *L2 classification* | | | | |
| VGG16 vs ResNet152 with Places365 | 0.084 | 0.107 | 0.120 | **0.048** |
| VGG16 vs ResNet152 with ImageNet | **0.034** | 0.083 | 0.055 | 0.071 |
| VGG16 vs ResNet152 with OwnWeights | **0.050** | **0.009** | **0.030** | **0.005** |
| VGG16 with Places365 vs VGG16 with ImageNet | 0.560 | 0.689 | 0.913 | 0.495 |
| ResNet152 with Places365 vs VGG16 with ImageNet | 0.998 | 0.915 | 0.954 | 0.960 |

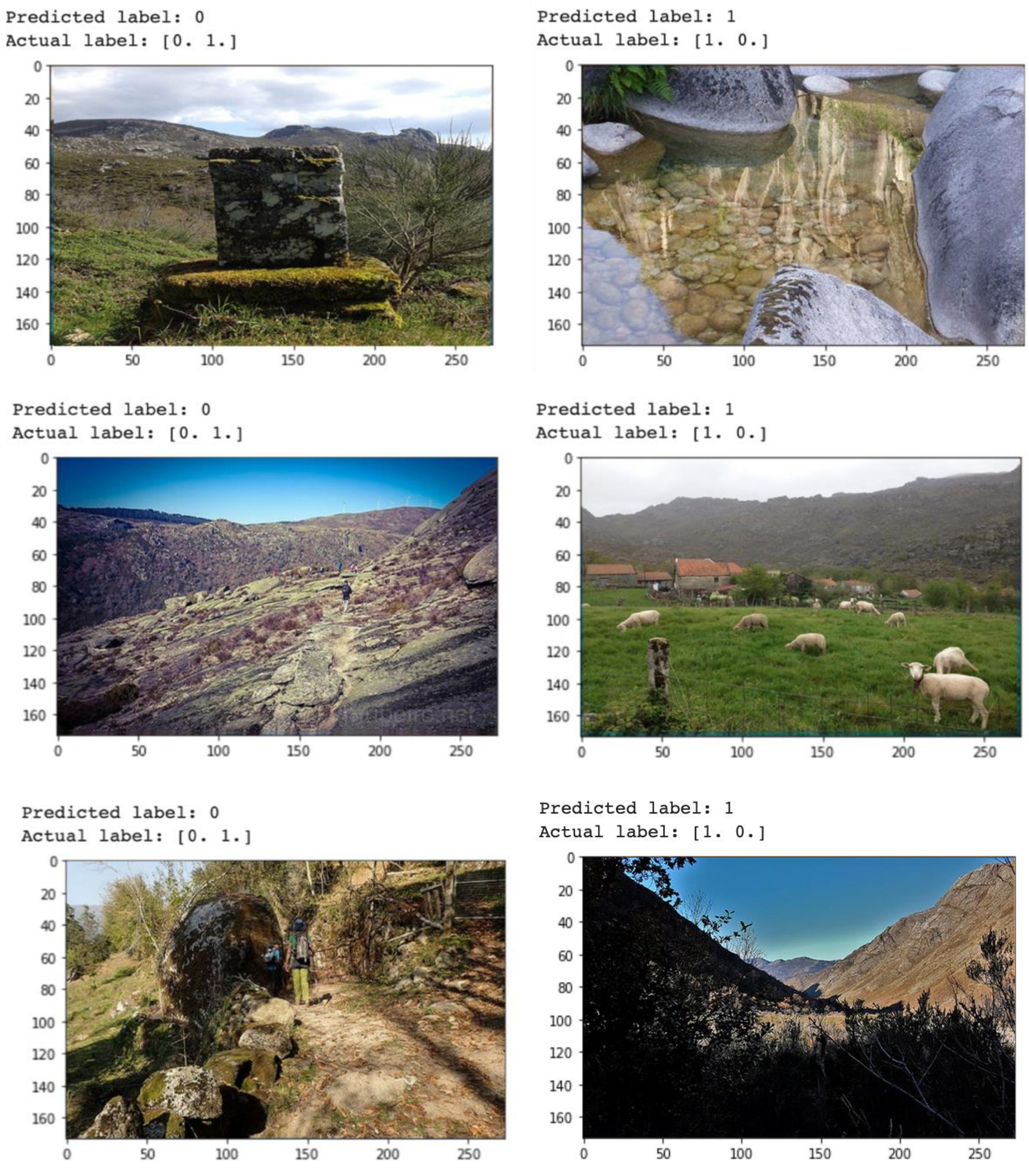


**Fig. S1.** Examples of images where the classes were swapped by the ResNet152 model with Places365 (first row), ImageNet (second row) and OwnWeights (third row). L1 classification of Peneda-Gerês: 0 – Nature, 1 – Human.

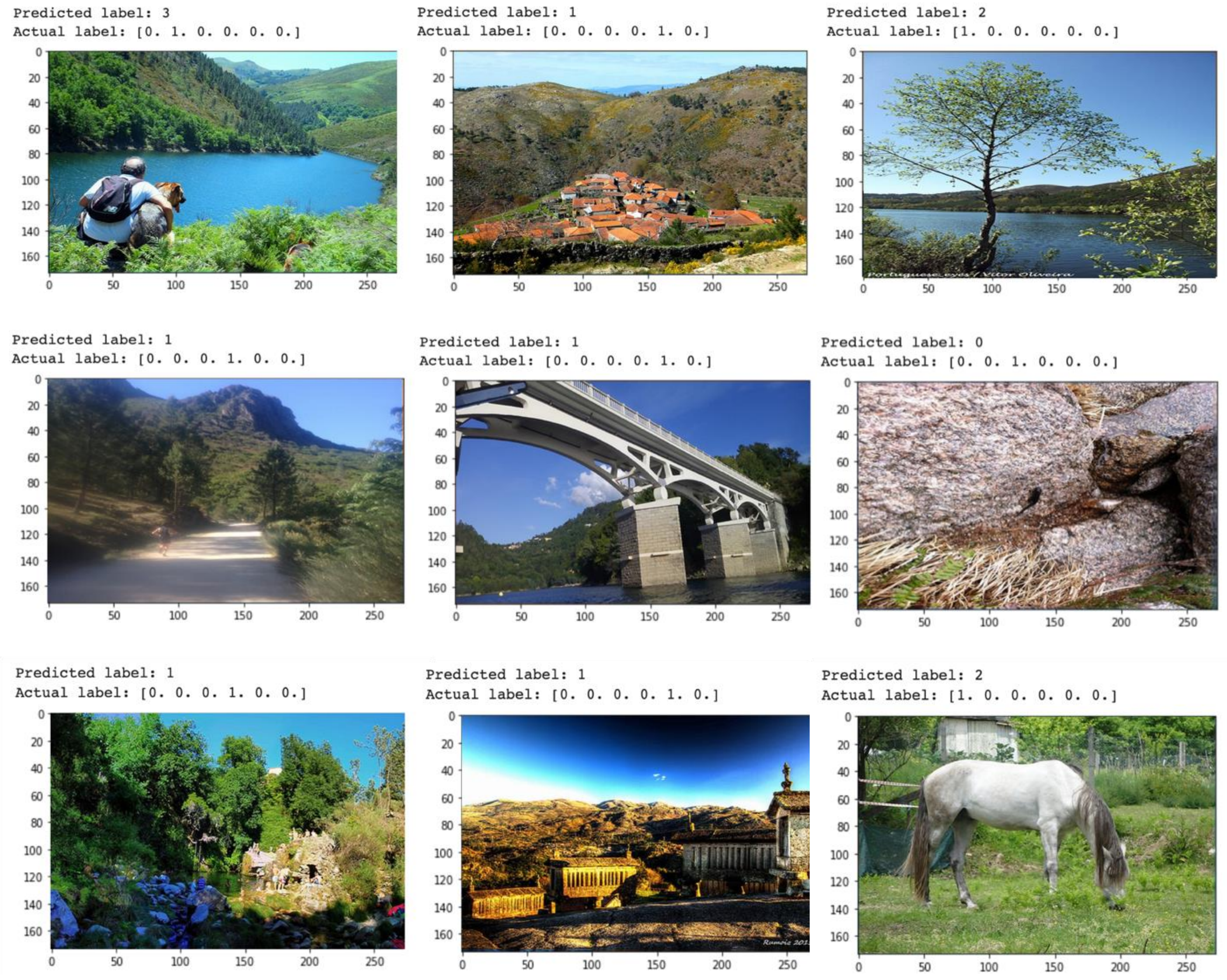


**Fig. S2.** Examples of images where the classes were swapped by the ResNet152 model with Places365 (first row), ImageNet (second row) and OwnWeights (third row). L2 classification of Peneda-Gerês: 0 – Species, 1 – Landscape, 2 – Nature, 3 – Human activities, 4 – Human structures, 5 – Posing.

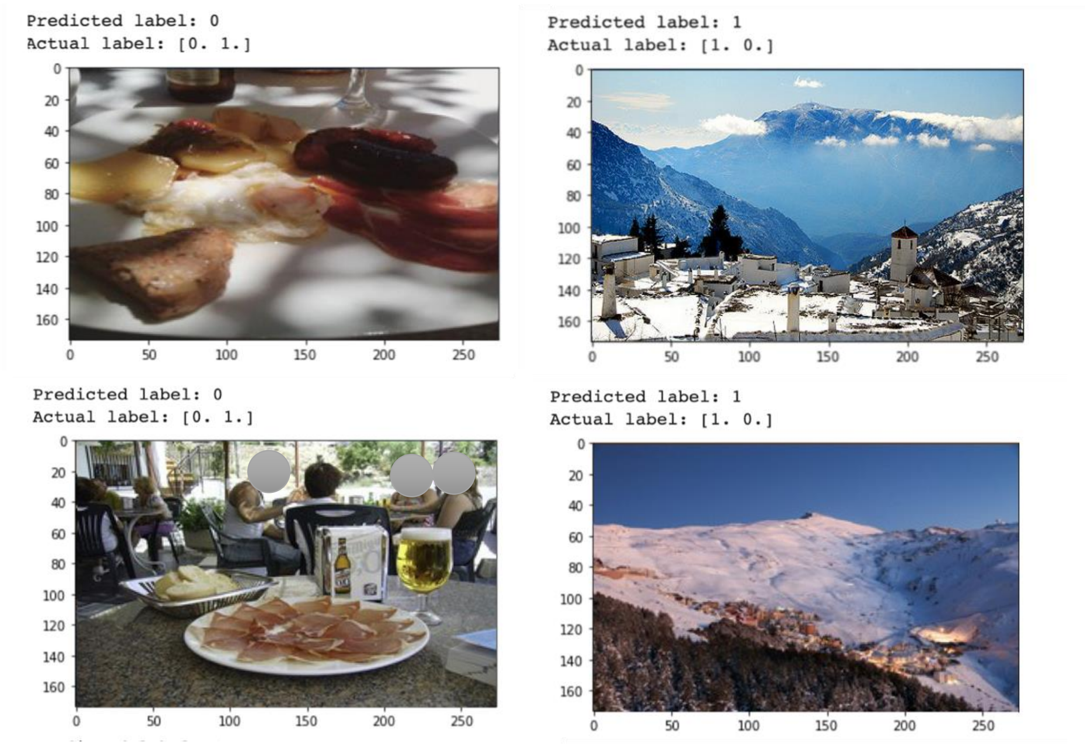


**Fig. S3.** Examples of images where the classes were swapped by ResNet152 model with ImageNet (first row) and OwnWeights (second row). L1 classification of Sierra Nevada: 0 – Nature, 1 – Human.

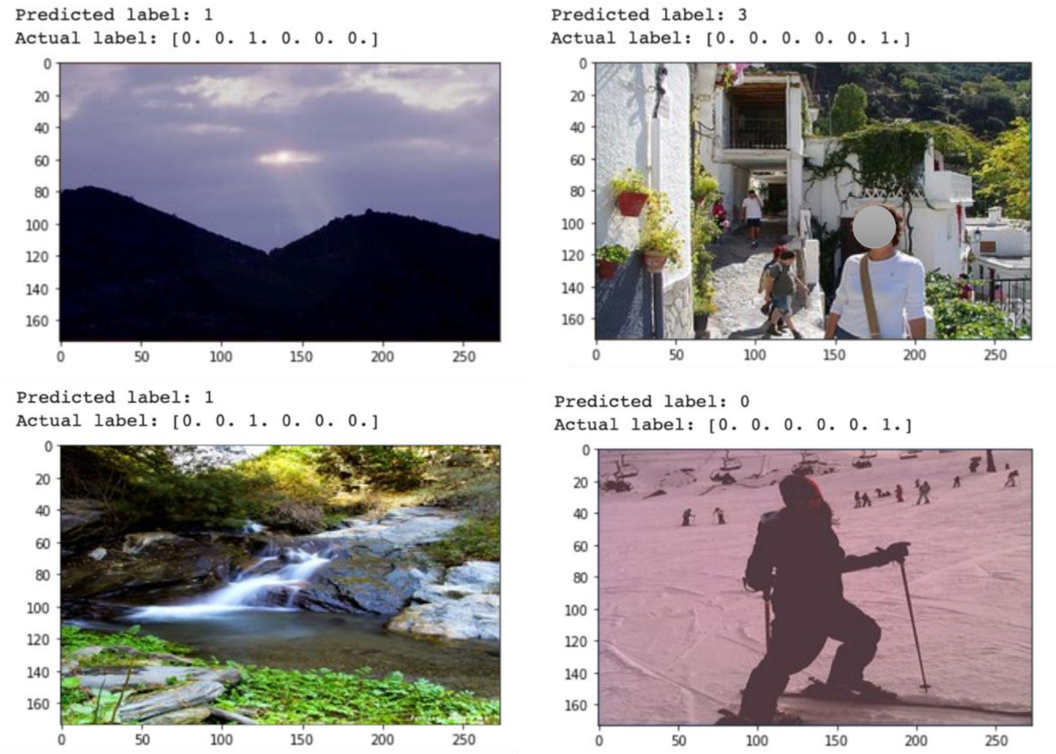


**Fig. S4.** Examples of images with classes for which the algorithm most failed the classification in ResNet152 with ImageNet (first row) and OwnWeights (second row). L2 classification of Sierra Nevada: 0 – Species, 1 – Landscape, 2 – Nature, 3 – Human activities, 4 – Human structures, 5 – Posing.
